# Analysis of the heterogenous structural states of the hexameric ATPase PilU of the Type IV pili from *Vibrio cholerae*

**DOI:** 10.64898/2026.02.06.704419

**Authors:** Yirui Guo, Shantanu Shukla, George Minasov, Nicole L. Inniss, Thomas Klose, Valerie L. Tokars, Alfonso Mondragón, Zbyszek Otwinowski, Dominika Borek, Karla J. F. Satchell

## Abstract

Type IV pili (T4P) mediate surface motility, host interactions, and DNA uptake through cycles of extension and retraction. While the primary retraction ATPase PilT has been extensively characterized, its homolog PilU remains less well understood despite being demonstrated as a PilT-dependent retraction ATPase. Here, we determined six PilU structures by cryo-electron microscopy and X-ray crystallography. The structures reveal a homohexameric assembly stabilized by interactions between the C-terminal catalytic domain of one subunit and the N-terminal PAS-like domain of a neighboring subunit. PilU adopts multiple conformational states, exhibiting different combinations of open and closed interfaces even in the absence of nucleotide. Comparison with PilT highlights structural features that likely underlie PilU’s weak ATPase activity and its dependence on PilT for function. Together, these findings provide a structural framework for understanding PilU’s role within the T4P retraction machinery.

## Introduction

Type IV pili (T4P) are dynamic, filamentous surface appendages found in many bacteria and archaea, where they play key roles in adhesion, motility, biofilm formation, DNA transformation, and host interaction (Craig et al., 2019; Singh et al., 2022). These supramolecular assemblies are composed primarily of pilin subunits assembled into a helical fiber via a conserved secretion system shared with type II secretion machineries (Gold et al., 2015; Karuppiah et al., 2016). Powered by dedicated cytoplasmic ATPases, T4P can rapidly assemble and extend from the bacterium. In pathogenic species, such as *Neisseria gonorrhoeae*, *Pseudomonas aeruginosa*, and *Vibrio cholerae*, a subclass of Type IV pili known as the Type IVa pili (T4aP) are also able to reverse the assembly process to retract, a process critical for host colonization, immuno-evasion, surface adhesion, and DNA uptake (Craig et al., 2004). In *V. cholerae*, two distinctive T4aP can be assembled. The extended mannose-sensitive hemagglutination (MSHA) pilus is a T4aP required for initial surface attachment with pilus retraction facilitating surface adhesion and initiation of biofilm formation (Watnick et al., 1999; Floyd et al., 2020). The second T4aP is not required for cell adherence or biofilm formation (Fullner and Mekalanos, 1999; Watnick et al., 1999), but instead is required for chitin-stimulated competence and DNA transformation (Meibom et al., 2005; Seitz and Blokesch, 2013), in which free DNA in the environment binds the tip of an extended pilus and is subsequently brought into the bacterium by pilus retraction for integration into the chromosome (Ellison et al., 2018). Due to their structural complexity, dynamic behavior, and evolutionary conservation, a detailed understanding of T4aP extension and retraction systems is crucial for elucidating the molecular mechanisms underlying bacterial adaptation to distinct environment (Craig et al., 2019; Ligthart et al., 2020).

During T4aP extension and retraction, pilin subunits in the periplasmic space are assembled into the elongating pilus through the coordinated action of the PilC-homolog inner-membrane platform proteins and the cytoplasmic motor ATPases (LaPointe and Taylor, 2000; Craig et al., 2019). The platform proteins serve as anchors for the ATPases and as a conduit for rotation of the pilus platform, enabling the addition of pilin subunits to the base of the extending pilus with the help of extension ATPases such as PilB and MshE. In contrast, the alternative motor ATPase PilT energizes pilus rotation in the reverse direction, during which pilin subunits are removed from the base of the retracting pilus and subsequently degraded (Chiang et al., 2008; Reindl et al., 2013; Chang et al., 2016; Chang et al., 2017).

The extension ATPase PilB for the competence pilus and MshE for the MSHA pilus as well as the shared retraction ATPase PilT have been extensively investigated through biochemical and structural studies of pili from several bacterial species (Craig et al., 2019). Despite driving opposing reactions, the two ATPases share a conserved architecture, assembling as hexameric rings composed of repeating subunits (Satyshur et al., 2007; Misic et al., 2010; Mancl et al., 2016; McCallum et al., 2017; McCallum et al., 2019). Each subunit contains a Per/Arnt/Sim (PAS)-like domain at the N-terminus (NTD) and a RecA-like motor domain at the C-terminus (CTD). The hexameric ring undergoes large-scale conformational changes associated with ATP binding and hydrolysis. These nucleotide-dependent rearrangements are thought to generate rotational forces that act through the PilC/MshG platform proteins, with PilB/MshE and PilT likely inducing rotation in opposite directions.

Some Type IV pilus genetic operons, including for the *Vibrio* competence and MSHA pili, carry genes that encode a third ATPase called PilU, a homolog of the retraction ATPase PilT. PilU and PilT share over 40% amino acid sequence identity (Figure S1). Several functional studies have shown that PilU acts as a PilT-dependent retraction ATPase, promoting pilus retraction only in the presence of PilT. Notably, this requirement is independent of PilT ATPase activity, as PilU-mediated retraction occurs even when PilT is catalytically inactive, in both competence and MSHA pili (Adams et al., 2019; Chlebek et al., 2019; Floyd et al., 2020). However, in some systems, both catalytically active PilT and PilU are required for twitching motility (Barnshaw et al., 2025). Additionally, it has been demonstrated that PilU can interact directly with PilT and PilC (Chlebek et al., 2019).

Despite these biochemical, genetic and structural modeling evidence suggesting that PilU contributes to efficient retraction in a PilT-dependent manner(Adams et al., 2019; Chlebek et al., 2019; Teipen et al., 2025), only one crystal structure of PilU from *Pseudomonas aeruginosa* (PDB: 9N32, resolution 4.54 Å) has been available to date (Barnshaw et al., 2025). In this work, we present a series of PilU structures determined by cryogenic electron microscopy (cryo-EM) and X-ray crystallography, providing insights into its architecture, nucleotide state, and potential mechanistic relationship with PilT and the T4aP system.

## Methods

### Cloning of PilU

PilU residues 2-368 from *V. cholerae* El Tor E7946 (ATQ45583.1) were cloned into a pET 15b vector with a 6×His N-terminal affinity tag MAGGSGGHHHHHHAGGAGG as previously described (Chlebek et al., 2019). The plasmid was kindly provided by Ankur Dalia (Indiana University).

### Purification of PilU

Recombinant PilU was expressed in *Escherichia coli* BL21(DE3)-Magic cell in 3 liters of Luria-Bertani media supplemented with 200 μg/ml ampicillin and 50 μg/ml kanamycin at 25 °C. The bacterial pellets collected by centrifugation were resuspended in 10 mM Tris-HCl pH 8.3, 0.5 M NaCl, 10 % glycerol, 0.1 % IGEPAL CA-630, flash frozen, and stored at −30 °C until purification. Thawed cells were lysed by sonication and lysate was clarified by centrifugation. The protein was purified using an ÅKTAxpress system (GE Healthcare) as previously described with some modifications (Shuvalova, 2014). The supernatant was loaded onto a HisTrapFF (GE Healthcare) column in loading buffer (10 mM Tris-HCl pH 8.3, 500 mM NaCl, 1 mM Tris (2-carboxyethyl) phosphine (TCEP), and 5 % glycerol). The column was washed with 10 column volumes (cv) of loading buffer and 10 cv of washing buffer (10 mM Tris-HCl pH 8.3, 1M NaCl, 25 mM imidazole, 5 % glycerol). The protein was eluted with elution buffer (10 mM Tris pH 8.3, 500 mM NaCl, 1 M imidazole) and then loaded onto a Superdex 200 26/600 column and separated in loading buffer. Samples for cryo-EM were concentrated to 10 mg/ml and used immediately. Samples for crystallization were flash-frozen in 20 % glycerol and stored at -80 °C until use.

### Cryo-EM grid preparation and data collection

The purified samples were applied to Quantifoil R 1.2/1.3 400 mesh copper grids. The grids were glow discharged for 60 s at 25 mA with a PELCO easiGlow™ Glow Discharge Cleaning System to obtain a hydrophilic surface. The glow-discharged grids were used to prepare vitrified samples with the Thermo Scientific Vitrobot Mark IV System. ATP or ADP were added to the protein sample at a final concentration of 3 mM and incubated for 5 min on ice before grid preparation. We applied 4 µl of purified protein to the glow-discharged surface of the grid at 4 °C at 100 % humidity with a wait time of 60 s, blot force of either 6 - 7 and blot time of 5 - 6 s.

The data were acquired using a 300 kV Titan Krios G1 microscope (Thermo Fisher) equipped with a K3 Summit detector and a BioQuantum K3 energy filter run in super-resolution mode at a nominal magnification of 130,000×, with a physical pixel size of 1.078 Å. A phase plate was not used, and the objective aperture was not inserted. Leginon was used for automated data collection in beam-image shift mode with 9 images collected per stage movement, over a defocus range from -0.5 µm to -2.5 µm and with beam-image shift compensation (Suloway et al., 2005; Cheng et al., 2021). The slit width of the Energy Filter was set to 20 eV. Movies were dose-fractionated into 40 frames with a total dose of ∼50 e^−^/Å^2^.

### Cryo-EM data processing

Three batches of data (dataset 1: apo, 3,559 movies; dataset 2: in the presence of ATP, 3,788 movies; dataset 3: in the presence of ADP, 3,836 movies) were collected and processed independently. All movies were imported into CryoSPARC v4.2-4.4 followed by patch motion correction using binning of 2. The contrast transfer function (CTF) was then estimated using patch CTF estimation (Punjani et al., 2017). Particles were first picked using the blob picker with a diameter range of 70-140 Å (2,793,135 particles for dataset 1; 2,668,741 particles for dataset 2; 1,967,136 particles for dataset 3). Picked particles were extracted with a box size of 432 pixels and cleaned by several rounds of 2D classifications. Classes showing clear protein features were kept for 3D reconstruction (356,413 particles for dataset 1; 366,165 particles for dataset 2; 274,583 particles for dataset 3).

Due to high heterogeneity observed in the particles, standard multi-class *ab initio* reconstruction followed by heterogeneous refinement approach did not result in maps with resolvable secondary structure features. As a result, multiple 3D classifications on different sets of particles were performed to capture particles with distinct conformations.

Particles from dataset 1 (356,413) were sent to an *ab initio* reconstruction job with one class followed by 3D classification with 10 total classes, a target resolution of 6 Å and an initial learning rate of 0.2. Particles were classified into six final classes. Homogeneous refinement was carried out on all classes and only one class (class 0: 101,834 particles) yielded a map with resolvable secondary structure features. After removing duplicate particles and repeating homogenous refinement, PilU form 1 is resolved to 3.44 Å in C3 symmetry with 93,306 particles (Figure S2, S3).

Inspection of the 2D class averages showed that ATP or ADP addition did not significantly alter the distribution of major views, overall particle shape or apparent symmetry. As a result, particles from datasets 1, 2 and 3 were combined for further processing. After removing duplicates, 860,736 pooled particles were subjected to *ab initio* reconstruction to create the consensus volume for several downstream 3D classification tasks.

The first 3D classification was performed on the 860,736 particles with 10 total classes, a target resolution of 6 Å and an initial learning rate of 0.4, yielding 5 final classes. After removing particles that resemble form 1 (class 0), 3D classification with the same parameters was performed again on the remaining 661,407 particles, which resulted in 5 final classes. Followed by homogeneous refinement for each class, PilU form 2 (class 2) was resolved to 3.68 Å in C2 symmetry with 114,357 final particles (Figure S2, S3).

To further resolve distinct conformations of PilU, an additional 3D classification was performed on the 860,736 particles using 10 total classes, a target resolution of 6 Å and an initial learning rate of 0.3, yielding 7 final classes. PilU forms 3 and 4 were resolved by homogeneous refinement to 3.49 Å (class 1) and 3.76 Å (class 2) in C2 symmetry, with 144,012 and 128,808 final particles, respectively (Figure S2, S3).

Local resolutions were estimated using the local resolution estimation job in cryoSPARC (Figure S3). Directional resolution anisotropy was assessed by 3D FSC analysis using the 3DFSC software package (Tan et al., 2017; Aiyer et al., 2021). The near-spherical FSC distributions indicate minimal preferred particle orientation and limited resolution anisotropy for all four reconstructions (Figure S4).

To visualize the motion of the hexameric ring, 3D Variability analysis was conducted on the clean particles from dataset 1 (Punjani and Fleet, 2021). Three modes were resolved in the analysis with C1 symmetry and filter resolution of 6 Å. Results were exported using 3D Variability Display job (simple mode) with 20 frames per mode. Movies for each mode/component were generated by ChimeraX (Goddard et al., 2018; Pettersen et al., 2021; Meng et al., 2023).

### Cryo-EM model building

Initial atomic models were obtained by docking an Alphafold2 model of PilU to the maps using Molrep (Vagin and Teplyakov, 2010; Jumper et al., 2021). Maps were also processed by *Phenix.resolve_cryo_em* for presentation and manual model building (Terwilliger et al., 2020). The model was iteratively rebuilt manually in Coot on the density modified maps and refined using *Phenix.real_space_refine* and Servalcat on raw maps (Emsley and Cowtan, 2004; Krissinel and Henrick, 2004; Emsley et al., 2010; Murshudov et al., 2011; Brown et al., 2015; Afonine et al., 2018; Liebschner et al., 2019; Yamashita et al., 2021). Model statistics are presented in Table 1. The atomic coordinates (PDB: 10MY, 10MZ, 10NA and 10NB) and maps (EMD-75296, EMD-75297, EMD-75298 and EMD-75299) have been deposited in the Protein Data Bank (http://wwpdb.org/) and the Electron Microscopy Data Bank (https://www.ebi.ac.uk/emdb/), respectively.

**Table 1.**
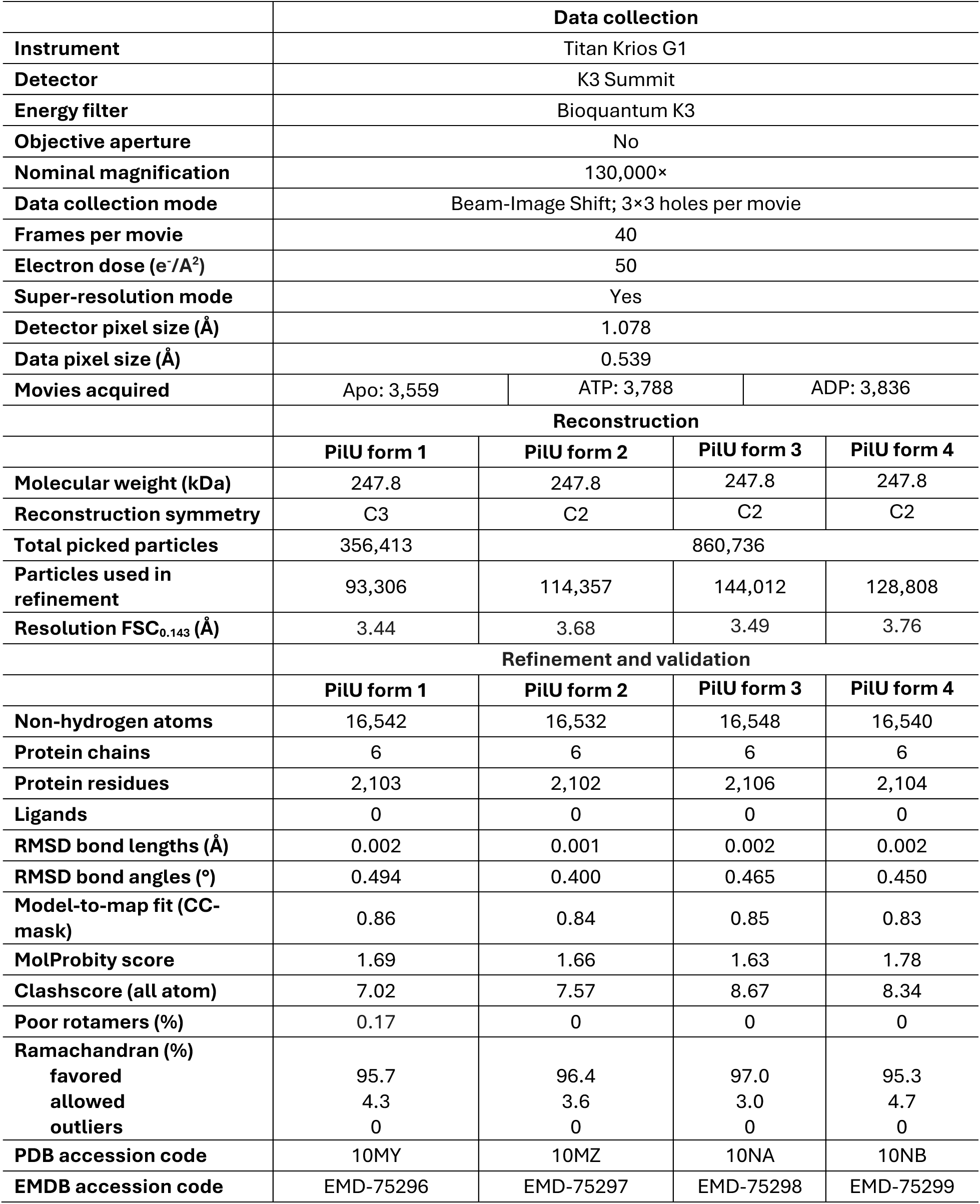
Cryo-EM data collection, processing and model building.

### Crystallization of PilU

Purified PilU was thawed and dialyzed into 10 mM Tris-HCl pH 8.3, 500 mM NaCl and 1 mM TCEP and concentrated to 15 mg/ml prior to crystallization trials. ATP or ADP were added to PilU at a final concentration of 1-3 mM and incubated for 30 minutes. Mixtures were spun at 12,000 RPM, 4 °C for 10 minutes before setting up 2 μl crystallization drops (1 μl protein: 1 μl reservoir solution) in 96-well crystallization plates (Corning) using commercial AmSO4, JCSG+, Classics II, PACT, PEG’s and PEG’s II (QIAGEN) crystallization screens. Data set #1 was collected from a crystal grown in the presence of 1 mM ATP from the condition containing 0.2 M sodium nitrate, 0.1 M Bis Tris propane pH 8.5, 20 % (w/v) PEG 3350 (PACT, #89), which was cryoprotected using well solution. Data set #2 was collected from the crystal grown in the presence of 3 mM ADP from the condition containing 0.2 M cadmium sulfate and 2.2 M ammonium sulfate (AmSO4, #13), which was cryoprotected using 2 M lithium sulfate. Crystals were flash frozen in liquid nitrogen for data collection.

### X-ray crystal structure determination

Each dataset was collected from a single crystal at beam line 21ID-D (crystal form 1, PilU form 6) and 21ID-F (crystal form 2, PilU form 5) of the Life Sciences-Collaborative Access Team (LS-CAT) at the Advanced Photon Source (APS), Argonne National Laboratory. Images were indexed, integrated, and scaled using HKL-3000 (Otwinowski and Minor, 1997; Minor et al., 2006). Data quality, structure refinement and the final model statistics are shown in Table 2.

**Table 2.** X-ray crystallography data processing and model building.

|  | **Crystal form 1 (PiIU form 6)** | **Crystal form 2 (PiIU form 5)** |
| --- | --- | --- |
| **Space group** | *P*2<sub>1</sub>2<sub>1</sub>2 | *I*222 |
| **Unit cell parameters (Å; °)** | a = 108.31, b = 133.73, c = 181.80;<br>$\alpha = 90.00, \beta = 90.00, \gamma = 90.00$ | a = 102.83, b = 120.83, c = 175.56;<br>$\alpha = 90.00, \beta = 90.00, \gamma = 90.00$ |
| **Resolution range (Å)** | 30.00 - 2.95 (3.00 - 2.95)<sup>a</sup> | 30.00 - 2.70 (2.75 - 2.70) |
| **No. of reflections** | 56,455 (2,794) | 30,441 (1,485) |
| ***R*<sub>pim</sub> (%)** | 4.4 (43.6) | 5.6 (53.7) |
| **Completeness (%)** | 99.3 (100.0) | 100.0 (100.0) |
| **$\langle I/\sigma(I) \rangle$** | 15.8 (1.6) | 16.1 (1.8) |
| **Multiplicity** | 11.9 (12.2) | 7.5 (7.5) |
| **Wilson *B* factor** | 71.6 | 41.6 |
| **Refinement and validation** |  |  |
|  | **Crystal form 1 (PiIU form 6)** | **Crystal form 2 (PiIU form 5)** |
| **Resolution range (Å)** | 29.57 - 2.95 (3.03 - 2.95) | 29.58 - 2.70 (2.77 - 2.70) |
| **Completeness (%)** | 99.2 (100.0) | 92.1 (63.0) |
| **No. of reflections** | 53,063 (4,091) | 26,612 (1,381) |
| ***R*<sub>work</sub>/*R*<sub>free</sub> (%)** | 22.3/26.5 (34.0/35.6) | 22.6/26.9 (28.8/31.8) |
| **Protein chains/atoms** | 6/16,526 | 3/8,230 |
| **Ligand/Solvent atoms** | 0/86 | 275/106 |
| **Mean temperature factor (Å<sup>2</sup>)** | 92.0 | 50.0 |
| **RMSD bond lengths (Å)** | 0.002 | 0.003 |
| **RMSD bond angles (°)** | 1.334 | 1.312 |
| **Ramachandran (%)** |  |  |
| **favored** | 96.0 | 97.0 |
| **allowed** | 4.0 | 3.0 |
| **outliers** | 0.0 | 0.0 |
| **PDB Accession Code** | 9Q6M | 9Q6W |

Crystal form 1 (PilU form 6) belongs to the orthorhombic space group *P2_1_2_1_2* with six protein chains in the asymmetric unit. The structure was determined by the molecular replacement method using the structure of PilT4 from *Geobacter metallireducens* (PDB: 6OJX) as a search model. The search model was split in two rigid bodies: the N-terminal domain (residues 4-98) and the C-terminal domain (residues 113-359), both were used to find a solution using PHASER (McCoy et al., 2007). The initial solution went through several rounds of refinement in REFMAC (Murshudov et al., 2011). Manual model corrections such as building missing residues and mutating the residues to match the sequence of PilU from *V. cholerae* were done using Coot (Emsley and Cowtan, 2004). The six chains in the asymmetric unit do not adopt a symmetric arrangement, consistent with cyclic symmetry 1. For comparison with cryo-EM reconstructions, we refer to it using Schoenflies notation as C1.

Crystal form 2 (PilU form 5) belongs to the orthorhombic space group *I222* with three protein chains in the asymmetric unit. The structure was determined by molecular replacement method, using the refined PilU structure from crystal form 1 as a search model in PHASER. The initial solution went through several rounds of refinement in REFMAC followed by manual model corrections in Coot. Because the complete hexamer is generated by application of crystallographic twofold symmetry to the three chains in the asymmetric unit, the assembly exhibits cyclic twofold symmetry. For comparison with cryo-EM reconstructions, this symmetry is described using Schoenflies notation as C2.

For both structures, the water molecules were generated automatically using ARP/wARP (Cohen et al., 2008) followed by additional rounds of refinement in REFMAC. Sulfate ions for crystal form 2 were fit into electron density maps manually. Translation–Libration–Screw (TLS) group corrections generated by the TLS Motion Determination (TLSMD) server (Painter and Merritt, 2006a; Painter and Merritt, 2006b) were used in the final round of REFMAC refinement for both structures. The quality of the model during refinement and the structure validation were performed using MolProbity (http://molprobity.biochem.duke.edu/) (Williams et al., 2018). The final structures were deposited to the Protein Data Bank (https://www.rcsb.org/) with the assigned PDB code 9Q6M for crystal form 1 (PilU form 6) and 9Q6W for crystal form 2 (PilU form 5).

### Multisequence alignment

Multisequence alignment was generated with PilU, PilT and PilB sequences from *V. cholerae*, *P. aeruginosa*, *Neisseria meningitidis* and *G. metallireducens* using the COBALT server (Papadopoulos and Agarwala, 2007) and visualization by Jalview (Clamp et al., 2004).

### Surface electrostatics analysis

Surface electrostatics analysis was performed using the APBS PyMOL plugin, with default settings corresponding to pH 7 (Jurrus et al., 2018), and PDB2PQR was used to add hydrogens and assign protonation states, atomic charges, and radii (Dolinsky et al., 2004).

### Binding energy calculation

Binding energy and dissociation constants between PilU domains and subunits at 25 °C were estimated using the PRODIGY server (Vangone and Bonvin, 2015; Xue et al., 2016).

### AlphaFold modeling

The AlphaFold models of PilU were calculated using the Google Colab platform AlphaFold2 (https://colab.research.google.com/github/sokrypton/ColabFold/blob/main/beta/AlphaFold2_adv anced.ipynb) (Jumper et al., 2021) with default settings (num_models = 5, ptm option, num_ensemble = 1, max_cycles = 3, num_relax = 0, tol = 0, num_samples = 1). The model with the highest pTMscore was used as a template for molecular replacement in atomic model building into cryo-EM maps. The AlphaFold model of PilU/PilT complex was generated using AlphaFold3 server (https://alphafoldserver.com/) with default parameters (Abramson et al., 2024), yielding an ipTM score of 0.65 and a pTM score of 0.67.

## Results

### PilU hexamer assumes multiple conformations

In this study, we solved four cryo-EM structures and two X-ray structures of PilU from *V. cholerae* strain E7946, all forming homohexameric rings with various conformations (Forms 1-6, Figure 1, Figure S2, S3, S4). These structures were solved in C1, C2 and C3 symmetry and the conformation variations were observed both in 2D classes (Figure S3a) and 3D reconstructions (Figure 1, Figure S2, S3f-m). For cryo-EM specifically, extensive 3D classification was utilized to separate the different forms of PilU (Figure S2). All of these observations suggest that PilU undergoes continuous motion similar to its close homolog PilT (McCallum et al., 2019).

**Figure 1.**
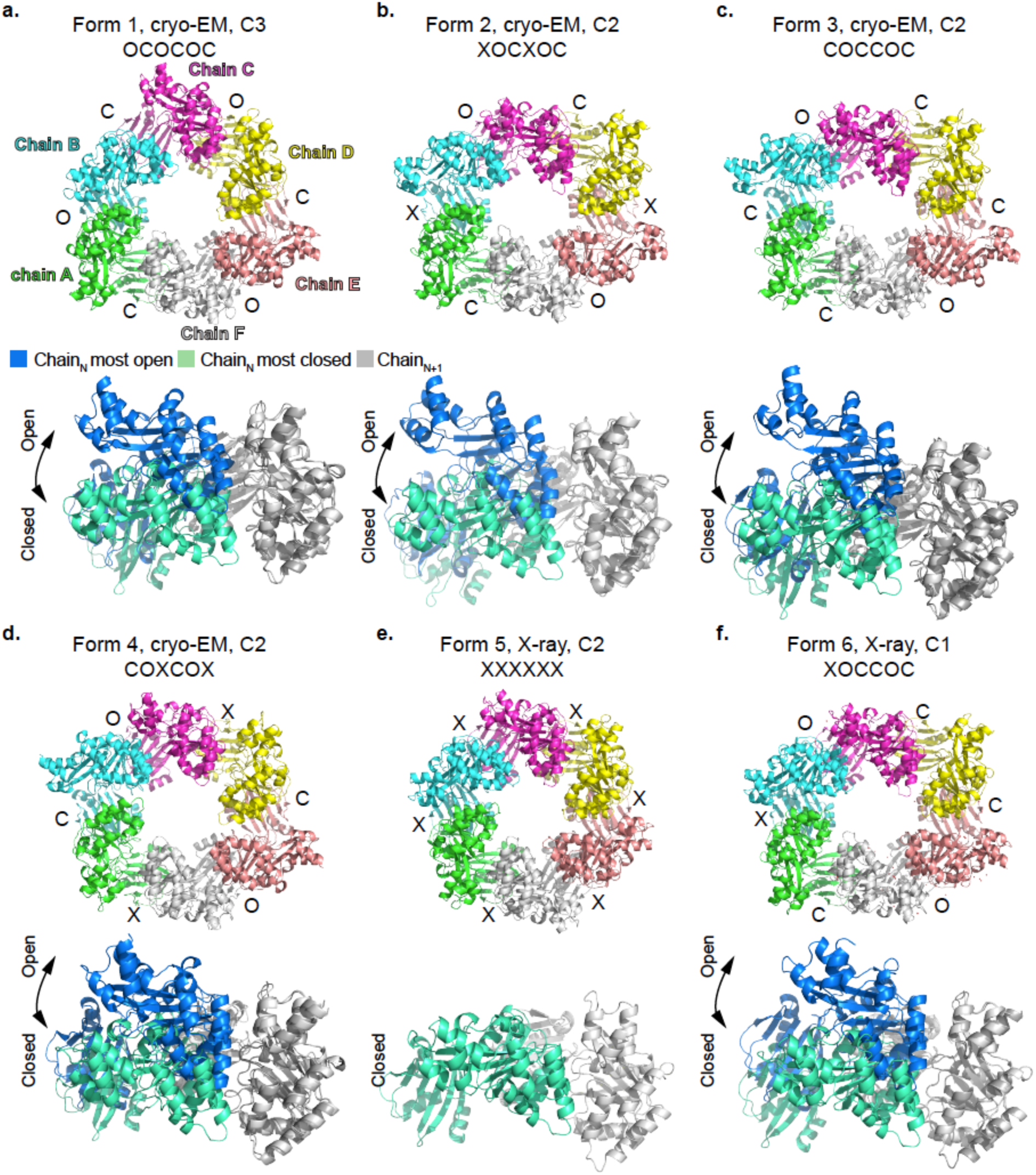
Four cryo-EM and two X-ray structures capture heterogeneous conformations of the PilU hexamers. Upper panel: PilU hexamer solved with different symmetry imposed, with each interface between adjacent subunits labeled as closed “C”, intermediate “X” or open “O”. Lower panel: the most closed interface and the most open interface from the same hexamer superimposed on CTD_N+1_ to show the largest conformational change within the hexamer. Form 5 only has interface resembles the closed conformation.

Each PilU subunit resembles the classic PilT/PilB fold with a PAS-like NTD and a catalytic CTD (Figure 2a, Figure S5). The CTD contains all the conserved motifs for ATP binding and hydrolysis. All subunits of the PilU forms 1-6 are in similar conformations with the Cα root-mean-squared-deviation (RMSD) in the range of 0.3 Å and 3.5 Å when superimposed using form 6 chain A as reference (Figure 2b). The upper end of this range is not due to conformational changes of the NTD or CTD individually, but arises from differences in the relative positioning of the two domains mediated by the flexible linker. When aligned by individual domains, Cα RMSD for NTD is in the range of 0.2 Å and 0.9 Å and Cα RMSD for CTD is in the range of 0.2 Å and 1.8 Å (Figure S6).

**Figure 2.**
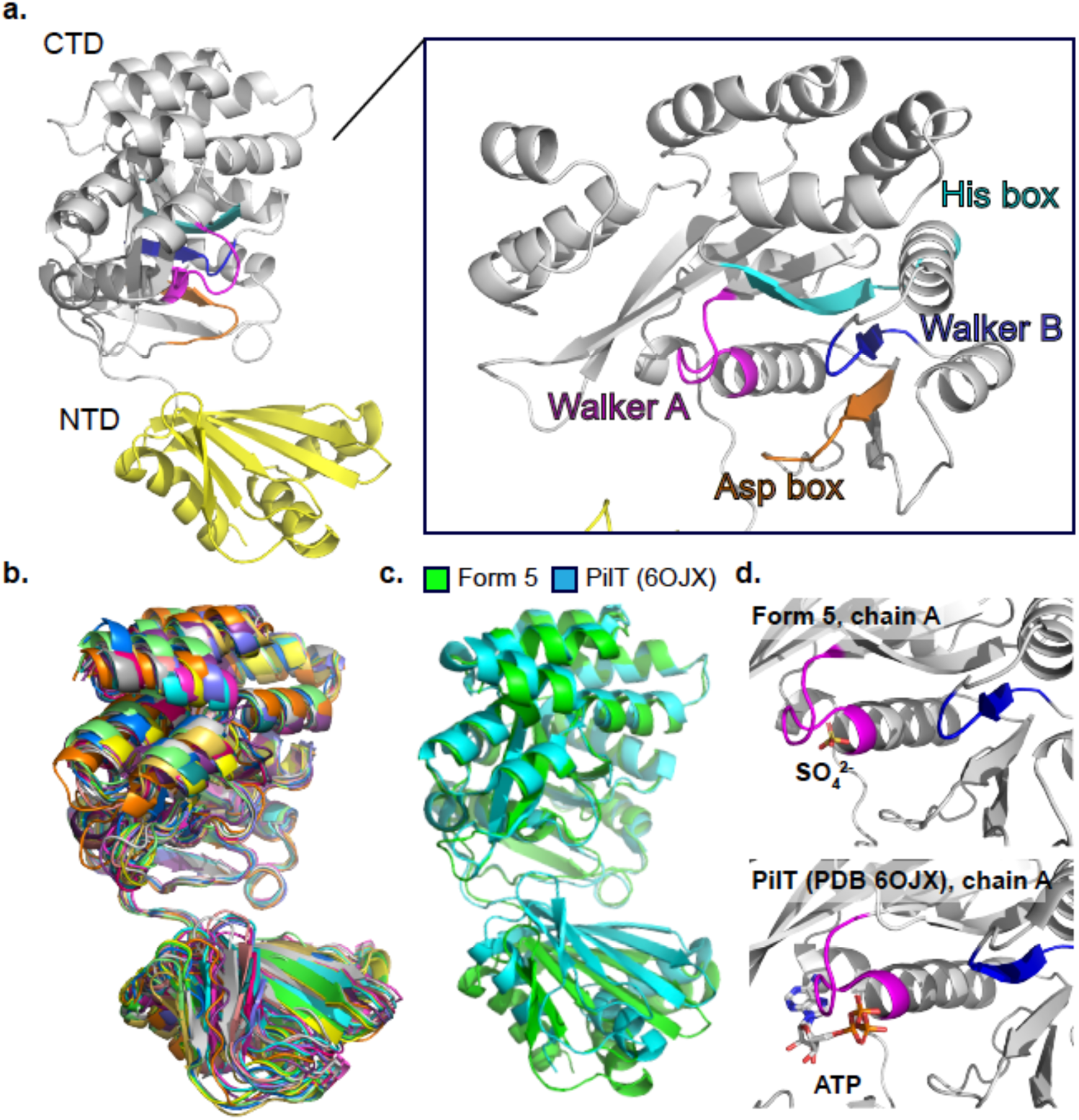
Structure of the PilU subunit. a) A PilU subunit consists of an NTD domain and a CTD domain. The catalytic pocket is located in CTD with each catalytic motif labeled (Walker A: magenta; Walker B: blue; his box: cyan; asp box: orange). b) Superimposition of PilU subunit from all six structures. c) Superimposition of the CTDs of PilU form 5 chain A and *G. metallireducens* PilT chain A (PDB: 6OJX). d) Comparison of the catalytic pockets between PilU and PilT.

To show a direct comparison between PilU and PilT, chain A of PilU form 5 was superimposed with chain A of *G. metallireducens* PilT (PDB: 6OJX), a structurally close homolog and a well-characterized retraction ATPase with clearly defined open and closed inter-subunit interfaces (Figure 2c). The global Cα RMSD between the two chains is 2.8 Å when aligned by both NTD and CTD. When only CTD is aligned, the Cα RMSD drops to 0.9 Å, as expected given the high sequence similarity. No nucleotide or Mg^2+^ density was observed in the structures, despite the addition of ATP and ADP in both the cryo-EM and crystallography samples. The only ligand modeled was a SO_4_ ion near the active site in the crystal structure of PilU form 5, whereas in its close homolog *G. metallireducens* PilT and *G. metallireducens* PilB, ATP or other nucleotide analogues were co-crystallized in the active sites (Figure 2d).

To form a ring structure, subunits of PilU are packed the same way as in other PilT and PilB structures where the CTD of subunit N (CTD_N_) is stacked on top of the NTD of subunit N+1 (NTD_N+1_) (Figure 3a). Following the method established by McCallum *et al*, the Cα distance between key residues on CTD_N_ and CTD_N+1_ (T218_N_-H227_N+1_ and T218_N_-L266_N+1_ in PilU; T211_N_-H230_N+1_, T211_N_-L269_N+1_ in PilT) are used to determine the open (O), intermediate (X) and closed (C) states of the interface between two adjacent PilU subunits (McCallum et al., 2019). A slightly relaxed distance cutoff was used in our analysis to avoid too many X states in the PilU structures but this relaxation did not change the results presented by McCallum *et al* in other PilT/VirB11-like family member structures. Briefly, if the Cα distance between T218_N_ and H227_N+1_ is larger than 12 Å and the Cα distance between T218_N_ and L266_N+1_ is less than 12 Å, the interface is open (O); if the Cα distance between T218_N_ and H227_N+1_ is less than 10.5 Å and the Cα distance between T218_N_ and L266_N+1_ is larger than 12 Å, the interface is closed (C); otherwise, the interface is in an intermediate state (X). With the above criteria, the status of the interface for each form is labeled in Figure 1 (Table 3). Superimposition of the CTD_N+1_ from the most open and closed interface within the same hexamer highlights the possible extreme position of CTD_N_ within each PilU conformation (Figure 1, Table 3). Each PilU homohexamer exhibited different degrees of open and closed states between subunits, with the most open interface found in form 3 between chain B and chain C, and the most closed interface found in form 6 between chain C and chain D, determined by the distance difference between T218_N_-H227_N+1_ and T218_N_-L266_N+1_. PilU form 1 resembles the crystal structure of *P. aeruginosa* PilU (PDB: 9N32, OCOCOC), with a Cα RMSD of 2.3 Å (Barnshaw et al., 2025). Based on distance measurements alone, 9N32 has a narrower closed interface (T220_N_-H229_N+1_ ∼8.6 Å and T220_N_-L268_N+1_ ∼16.3 Å) than PilU form 6 (T218_N_-H227_N+1_ ∼9.2 Å and T218_N_-L266_N+1_ ∼15.6 Å). However, because this interface in 9N32 is generated by crystallographic symmetry and the structure has limited side chain assignment due to low resolution, 9N32 was not included in the subsequent discussion of open and closed states.

**Figure 3.**
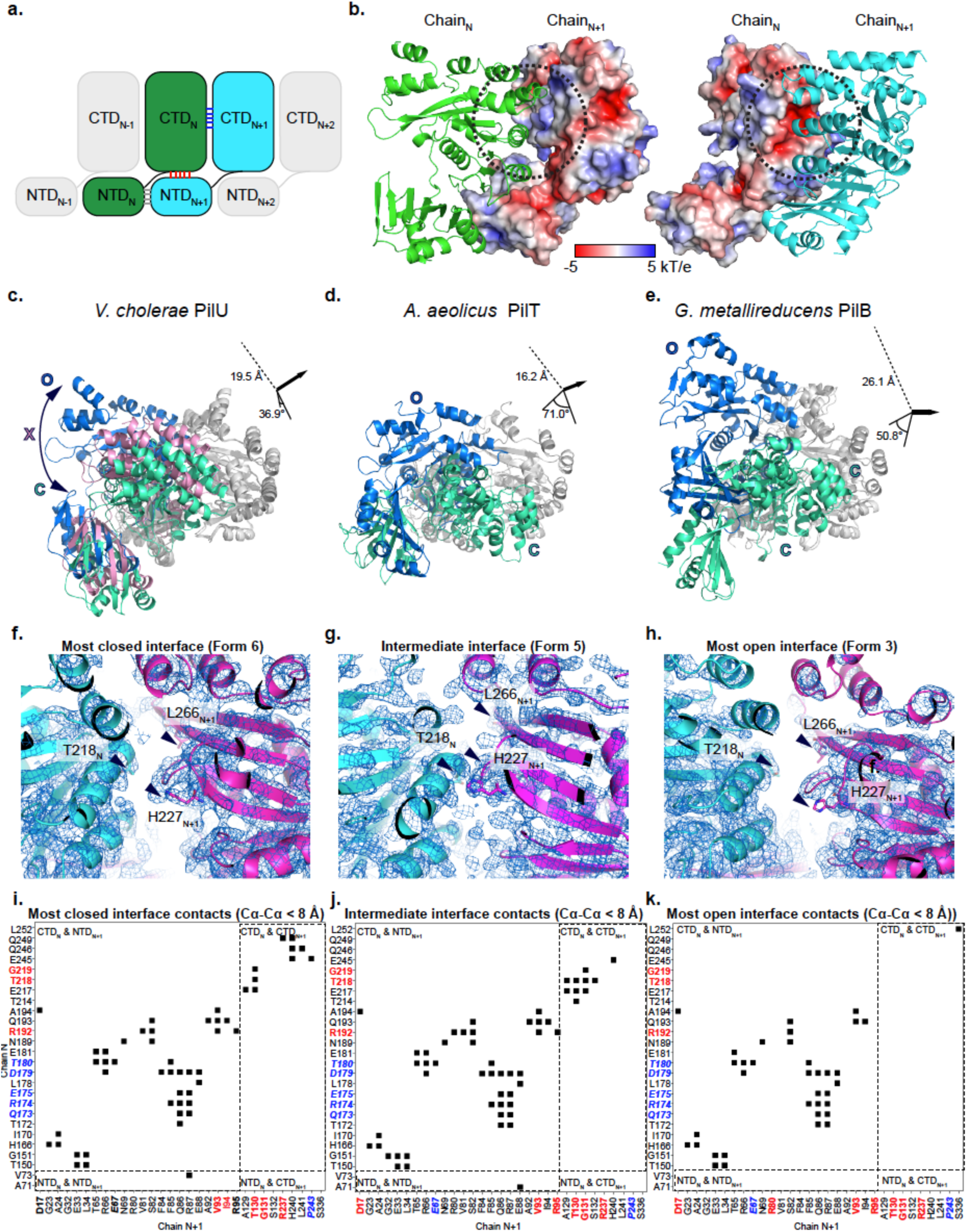
The closed and open interface of PilU. a) Schematic of subunit arrangement in PilU. b) Surface electrostatic potential of adjacent PilU subunits (PilU form 6, chains C and D). c) Conformational change between the most closed (green) and the most open (blue) interface across all PilU structures. An intermediate state is shown in pink. d) Conformational change between the most closed (green) and the most open (blue) interface across all PilT structures. e) Conformational change between the most closed (green) and the most open (blue) interface across all PilB structures. f) Density map of the most closed interface of PilU formed by chain C (cyan) and chain D (magenta) of form 6. g) Density map of the intermediate interface of PilU formed by chain A (cyan) and chain B (magenta) of form 5. h) Density map of the most open interface formed by chain B (cyan) and chain C (magenta) of form 3. i-k) Residues in contact in the most closed PilU interface, the intermediate PilU interface and the most open PilU interface, respectively. Contact is determined by residues with Cα-Cα distance less than 8 Å. Conserved residues across PilU, PilT and PilB are in bold red text. Conserved residues across PilU and PilT are in bold blue italics text.

**Table 3.** Open and closed interface determination between adjacent PilU subunits based on Cα distance T218_N_-H227_N+1_ and T218_N_ -L266_N+1_.

| PilU | Chains at interface | (1) Ca T218<sub>N</sub>-H227<sub>N+1</sub> (Å)* | (2) Ca T218<sub>N</sub>-L266<sub>N+1</sub> (Å)** | Distance difference (1) – (2) (Å) | O/C based on McCallum *et al* cutoff | O/C based on relaxed cutoff | Most-O/C within hexamer | Most-O/C across all hexamers |
| --- | --- | --- | --- | --- | --- | --- | --- | --- |
| **Form 1** | AB | 15.7 | 10.5 | 5.2 | O | O | Most-O | - |
|  | BC | 10.4 | 15.7 | -5.3 | X | C | Most-C | - |
| **Form 2** | AB | 12.9 | 14.8 | -1.9 | X | X | - | - |
|  | BC | 16.5 | 10.5 | 6 | O | O | Most-O | - |
|  | CD | 9.6 | 15.8 | -6.2 | X | C | Most-C | - |
| **Form 3** | AB | 9.7 | 15.6 | -5.9 | X | C | Most-C | - |
|  | BC | 18.9 | 11.2 | 7.7 | X | O | Most-O | Most-O |
|  | CD | 9.5 | 15.1 | -5.6 | X | C | - | - |
| **Form 4** | AB | 9.3 | 15.3 | -6 | X | C | Most-C | - |
|  | BC | 15.2 | 9.4 | 5.8 | O | O | Most-O | - |
|  | CD | 12.2 | 12.9 | -0.7 | X | X | - | - |
| **Form 5** | AB | 11.5 | 10.7 | 0.8 | X | X | Most-C | - |
|  | BC | 11.5 | 10.5 | 1 | X | X | - | - |
|  | CD | 11.6 | 9.8 | 1.8 | X | X | - | - |
| **Form 6** | AB | 10.3 | 12.2 | -1.9 | X | X | - | - |
|  | BC | 15.2 | 9.9 | 5.3 | O | O | Most-O | - |
|  | CD | 9.2 | 15.6 | -6.4 | X | C | Most-C | Most-C |
|  | DE | 9.9 | 15.3 | -5.4 | X | C | - | - |
|  | EF | 14.4 | 10.3 | 4.1 | O | O | - | - |
|  | FA | 9.3 | 15 | -5.7 | X | C | - | - |

\*Ca distance cutoff for open interface (O): T218<sub>N</sub> - H227<sub>N+1</sub> > 12 Å; T218<sub>N</sub> - L266<sub>N+1</sub> < 12 Å (<11 Å in McCallum *et al.*, 2019)
\*\*Ca distance cutoff for closed interface (C): T218<sub>N</sub> - H227<sub>N+1</sub> < 10.5 Å (<9 Å in McCallum *et al.*, 2019); T218<sub>N</sub> - L266<sub>N+1</sub> > 12 Å (>11 Å in McCallum *et al.*, 2019)

### Open and closed interfaces are created by significant domain rotation between adjacent subunits

Open, intermediate and closed interfaces are observed in all the PilU structures (Figure 1, Table 3). Since each subunit, when overlaid individually, did not show large conformational changes, the open and closed interfaces are primarily driven by the relative positions of adjacent subunits (N, N+1), which are bridged by residue contacts at several regions of each subunit. Based on the subunit arrangement within the hexameric ring, contacts can happen between NTD_N_ and NTD_N+1_, CTD_N_ and NTD_N+1_, and CTD_N_ and CTD_N+1_ (Figure 3a).

The surface of a PilU subunit exhibits an electrostatic potential between -5 and 5 kT/e (Figure 3b). In CTD_N_, α8 and α11 form a slightly negatively charged patch that lies in close proximity to a positively charged patch near α9 in CTD_N+1_. Helix α7 in CTD_N_ also forms a small positively charged patch that is in proximity with a negatively charged patch on β6 in NTD_N+1_ (Figure 3b, Figure S5). The slightly charged patches between adjacent CTDs likely guide interactions between subunits during conformational changes, while maintaining sufficient flexibility for the interface to switch between open and closed states without a high-energy barrier. This is consistent with the calculated binding free energies (ΔG) between CTD_N_ and CTD_N+1_, which range from -5.7 to -5.9 kcal/mol (K_d_ ∼ 50-60 μM) for the closed and open states. In contrast, the ΔG between CTD_N_ and NTD_N+1_ is approximately -10 kcal/mol (K_d_ ∼ 40 nM), providing the structural core that stabilizes the hexamer.

As previously mentioned, we have identified that the most closed state of all PilU structures is in form 6 between chains C and D, and the most open state of all PilU structures is in form 3 between chains B and C. To visualize the transition between closed and open conformations, form 6 chains C and D, and form 3 chains B and C were superimposed on CTD_N+1_ (using form 6 chain D as the reference). Form 5 chains A and B were also included to show the position of an intermediate state. To transition from the most closed state to the most open state, CTD_N_ must rotate by 37 ° and shift by 19.5 Å (Figure 3c). Similar movement was observed in PilT and PilB (McCallum et al., 2017; McCallum et al., 2019). The cases of the most extreme open/closed states of all PilT and PilB structures are found in *Aquifex aeolicus* PilT (PDB: 2GSZ) (Satyshur et al., 2007) and *G. metallireducens* PilB (PDB: 5TSH) (McCallum et al., 2017) (Figure 3de). In *A. aeolicus* PilT, a rotation of 71 ° and a translation of 16.2 Å of the CTD_N_ are required to transition between the most closed and most open states. In *G. metallireducens* PilB, a rotation of 50.8 ° and a translation of 26.1 Å are required. The movement observed in PilU between the most closed and most open states is modest compared to the extreme cases of PilT and PilB. In addition, the most open and closed states of PilT and PilB are observed in a single structure (e.g., the most closed and open states of *A. aeolicus* PilT are observed in PDB 2GSZ, the most closed and open states of *G. metallireducens* PilB are observed in PDB 5TSH), whereas in PilU the most closed and open states are found in separate structures (Form 3 for most open, and form 6 for most closed), suggesting that PilT and PilB can exhibit larger distortions of the hexameric ring (Figure 3c-e). Both *A. aeolicus* PilT (PDB: 2GSZ) and *G. metallireducens* PilB (PDB: 5TSH) structures contain six monomers in the asymmetric unit; therefore, the observed conformations are not imposed by crystallographic symmetry and likely represent biologically accessible states, although they may still be influenced by crystal packing. Nevertheless, due to limitations of the experimental methods, we may not have captured the full range of PilU conformations, and similarly extreme ring distortions may exist but were not observed in this study.

The contacting residues in the most closed, intermediate, and most open states of PilU were analyzed in the contact map using Cα distance < 8 Å as a cutoff to avoid influences from side chain conformation, as well as using PDBsum (Laskowski et al., 2018) considering side chains, to illustrate the shift of the subunit interface during the transition from the closed to the open state. The most closed state is represented by the interface between chains C and D in form 6, as indicated by the distance between T218_N_, H227_N+1_ and L266_N+1_ (Figure 3f). Residues in contacts in this closed state are found in between CTD_N_ and NTD_N+1_, between CTD_N_ and CTD_N+1_, and between NTD_N_ and NTD_N+1_. The types of contacts include hydrogen bonds, salt bridges and non-bounded interactions (hydrophobic and van der Waal’s). Major contacts between CTDs include regions around α8 in CTD_N_ (E217, T218, G219) and the loop between β7 and α6 in CTD_N+1_ (T130), as well as regions around α10, α11 (E245, Q246, Q249) in CTD_N_ and α9, α10 in CTD_N+1_ (R237, H240, L241, P243). Major contacts between CTD_N_ and NTD_N+1_ include β8-β10 in CTD_N_ and β1-3 in NTD_N+1_, between β9, α7 in CTD_N_ and α3, β5, β6 in NTD_N+1_ (Figure 3i, Figure S7a, Figure S8a-d). A special case is R192 in CTD_N_, which is making contacts with both CTD_N+1_ and NTD_N+1_ (Figure S7a, S8b). In the intermediate state (form 5, chains A, B), the interactions between CTD_N_ and NTD_N+1_ did not change much. However, contacts between α10, α11 in CTD_N_ (E245, Q246, Q249) and α9 in CTD_N+1_ (R237, H240, L241, P243) were reduced, and contacts between α8, β11 in CTD_N_ (T214, E217, T218, G219) and β7, α6 in CTD_N+1_ (A129, T130, G131, S132) were increased as CTD_N_ shifted away from the closed state (Figure 3gj, Figure S7b, Figure S8ef). The most open state is in form 3 between chains B and C, where the interactions between CTD_N_ and NTD_N+1_ are maintained, but the interactions between CTD_N_ and CTD_N+1_ are almost completely lost (Figure 3hk, Figure S7c).

### Proposed trajectory of hexameric ring movement based on structural similarities

To visualize the relationship among the six PilU conformations, we connected the structures based on pairwise minimum Cα RMSD, generating a putative trajectory that represents the most direct path linking all observed states. It should be noted that this trajectory may not necessarily match what happens in solution as the energy barrier to transition between open and closed states is low, making it very easy for the protein to revert to a previous state or sample multiple intermediate conformations rather than strictly following a single forward pathway.

To achieve best alignment across all PilU forms, the six structures are aligned pairwise with the CE algorithm (Shindyalov and Bourne, 1998). This algorithm enables optimal match between two structures regardless of chain ID. The Cα RMSD of the entire molecule (>1,800 atoms) is then calculated between each pair of structures and used as a proxy to determine the similarity between each form (low RMSD = more similar). The most probable path is the one that morphs through all structures with the smallest sum of RMSD.

From the pairwise overall RMSD measurement between PilU structures, it is clear that Forms 1 and 5 are the most distant from all other structures. Both form 1 and form 5 show a pairwise RMSD of ∼5.5-9 Å when compared to other PilU structures, while pairwise RMSD between form 2, 3, 4 and 6 are within the range of 1.6-4 Å (Table 4).

**Table 4.** Pairwise Cα RMSD between PilU structures.

|  | Form 1 | Form 2 | Form 3 | Form 4 | Form 5 | Form 6 |
| --- | --- | --- | --- | --- | --- | --- |
| Form 1 | - |  |  |  |  |  |
| Form 2 | 5.6 | - |  |  |  |  |
| Form 3 | 7.0 | 2.7 | - |  |  |  |
| Form 4 | 5.1 | 3.5 | 3.2 | - |  |  |
| Form 5 | 4.1 | 6.2 | 9.0 | 5.7 | - |  |
| Form 6 | 5.7 | 1.7 | 3.0 | 3.8 | 4.9 | - |

The best trajectory including all structures is then determined by finding the smallest sum of RMSD between all PilU forms. The shortest path to go through each structure is in the order of form 1, 4, 3, 2, 6, 5 (total path length sum of RMSD = 21.6 Å) (Figure 4, Movie S1). Considering that form 5 was not observed in the cryo-EM data both in 2D classes and 3D reconstruction, a trajectory using only forms 1,4,3,2,6 is also generated (Movie S2), and this trajectory is consistent with the result of 3D variability analysis (Movie S3). Form 5, although solved with C2 symmetry, exhibit features close to C6 symmetry. In addition, all interfaces between subunits in form 5 are denoted as X, which is unusual in comparison to all other PilU structures. As a result, form 5 could be a special conformation that is only stabilized by lattice contacts or crystallization conditions.

**Figure 4.**
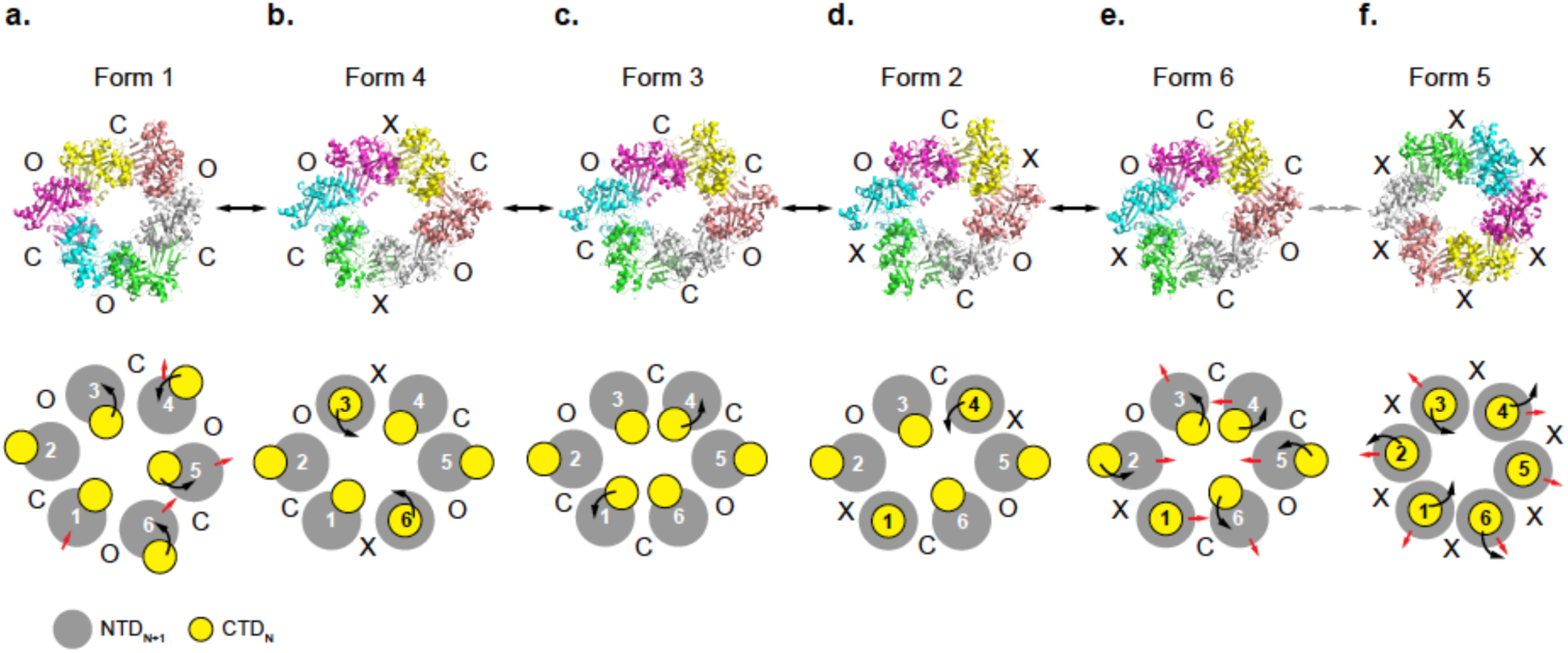
A potential conformational landscape connecting the observed states. A possible pathway of conformational transitions across all the structures was proposed based on the minimum RMSD between states. Black arrow represents the movement of the CTD, while red arrows represent global conformational changes of the hexameric ring.

Although this trajectory does not necessarily reflect the true conformational pathway in solution, it provides a qualitative framework for comparing structural rearrangements and identifying flexible regions within the complex. The sequence of models highlights progressive shifts in the CTD, suggesting how the protein may accommodate conformational changes during pilus retraction.

## Discussion

PilU ATPases are close homologs of the PilT retraction ATPases. The conformational heterogeneity of PilT is supported by both crystallographic and cryo-EM studies. Crystal structures have captured multiple conformations that correspond to C2-, C3-, and C6-equivalent symmetries, while several low-resolution cryo-EM maps, for which no atomic models were built, exhibit C2 and C3 symmetries (McCallum et al., 2019). Here we report similar observations of PilU by cryo-EM and crystallography. The structural similarity between PilU and PilT is expected given their sequence similarity. In PilT, ATP or ATP analogs promote closure of the inter-subunit interface, whereas ADP-bound PilT structures do not consistently correlate with specific open or closed states. This likely reflects the requirement to sequester and protect bound ATP within a closed interface until hydrolysis occurs, while ADP binding does not impose the same structural constraint. In this study, no nucleotide was observed in the active site, so all reported structures are considered to be in the apo state. Taken together with the observation that the same PilU conformations were found in both the apo and mixed datasets, and that PilU exhibits low ATPase activity *in vitro* (Chlebek et al., 2019), it is unlikely that significant ATP hydrolysis occurred under the conditions used. Thus, it is reasonable to assume that the conformations we resolved represent a series of ground-state conformations sampled by PilU until substrate (ATP) binding stabilizes a closed interface.

The arrangement of the closed (C), open (O), and intermediate (X) interface in the forms of OCOCOC, COCCOC, XXXXXX are found in both PilU and PilT structures, while OOOOOO, CCCCCC and OOCOOC conformations are only observed in PilT, and COXCOX, XOCXOC and XOCCOC are only observed in PilU. Among all PilT and PilU structures, OOCOOC is the most extreme conformation and was only observed in *A. aeolicus* PilT (PDB: 2GSZ). For PilU to adopt this conformation, larger rotations and translations of the CTD would be required.

In addition, compared to PilU, the gap between CTD_N_ and CTD_N+1_ in PilT structures is narrower in its closed state and wider in its open states, as demonstrated by the T218_N_-H227_N+1_ and T218_N_-L266_N+1_ Cα distances (Table 3). If the original distance cutoff by McCallum *et al* was applied, several closed and open states of PilU would need to be denoted as the intermediate state (Table 3). In other words, the open interface of PilT is more open than in PilU, and the closed interface of PilT is more closed than in PilU (McCallum et al., 2019). Based on this observation, we hypothesize that PilU alone does not exhibit strong ATPase activity because it lacks the proper arrangement of open and closed interfaces needed for ATP binding and effective catalysis. This is consistent with the weak ATPase activity measured for PilU *in vitro* (Chlebek et al., 2019). We propose that direct interaction with PilT is required for PilU to adopt the appropriate conformational states for ATP binding and hydrolysis, and to couple effectively to the rest of the T4aP machinery. In addition, it has been shown that apo PilT (PDB: 6OJY) can still adopt conformation similar to nucleotide bound PilT (PDB: 6OJZ), suggesting that the PilT’s overall conformation is not strictly determined by ATP binding or ATPase activity. This flexibility allows catalytically inactive PilT to adopt shapes that can stabilize a functional, catalytically competent conformation of PilU. Such an arrangement would enable pilus retraction to be powered by both PilT and PilU, or by PilU alone when PilT is inactive. Recent cryo-ET reconstruction of the *V. cholerae* competence pilus machine showed PilU-like density directly underneath PilT, providing evidence for direct PilT and PilU interaction *in vivo* (Maggi et al., 2025).

An AlphaFold model was generated to illustrate the possible interaction between PilT and PilU (Figure 5a-c). The predicted model is close to the CCCCCC form due to the C6 symmetry enforced by the algorithm (Abramson et al., 2024). Regardless of the state of the interface, the model predicts that the CTD of PilU interacts with two adjacent NTDs of PilT (Figure 5bc). Such interactions will limit PilU’s freedom of movement and tie its motion to PilT, facilitating synchronized movement driven by ATP hydrolysis of either one or both proteins (Figure 5d). When both PilT and PilU possess intact ATPase activity, the combined action effectively enhances the motor’s power output, enabling rapid pilus retraction, particularly under conditions that require strong retraction forces such as when pili are attached to a surface. Our modeling represents the same state compared to a similar modeling strategy reported as a preprint while this manuscript was in preparation (Teipen et al., 2025).

**Figure 5.**
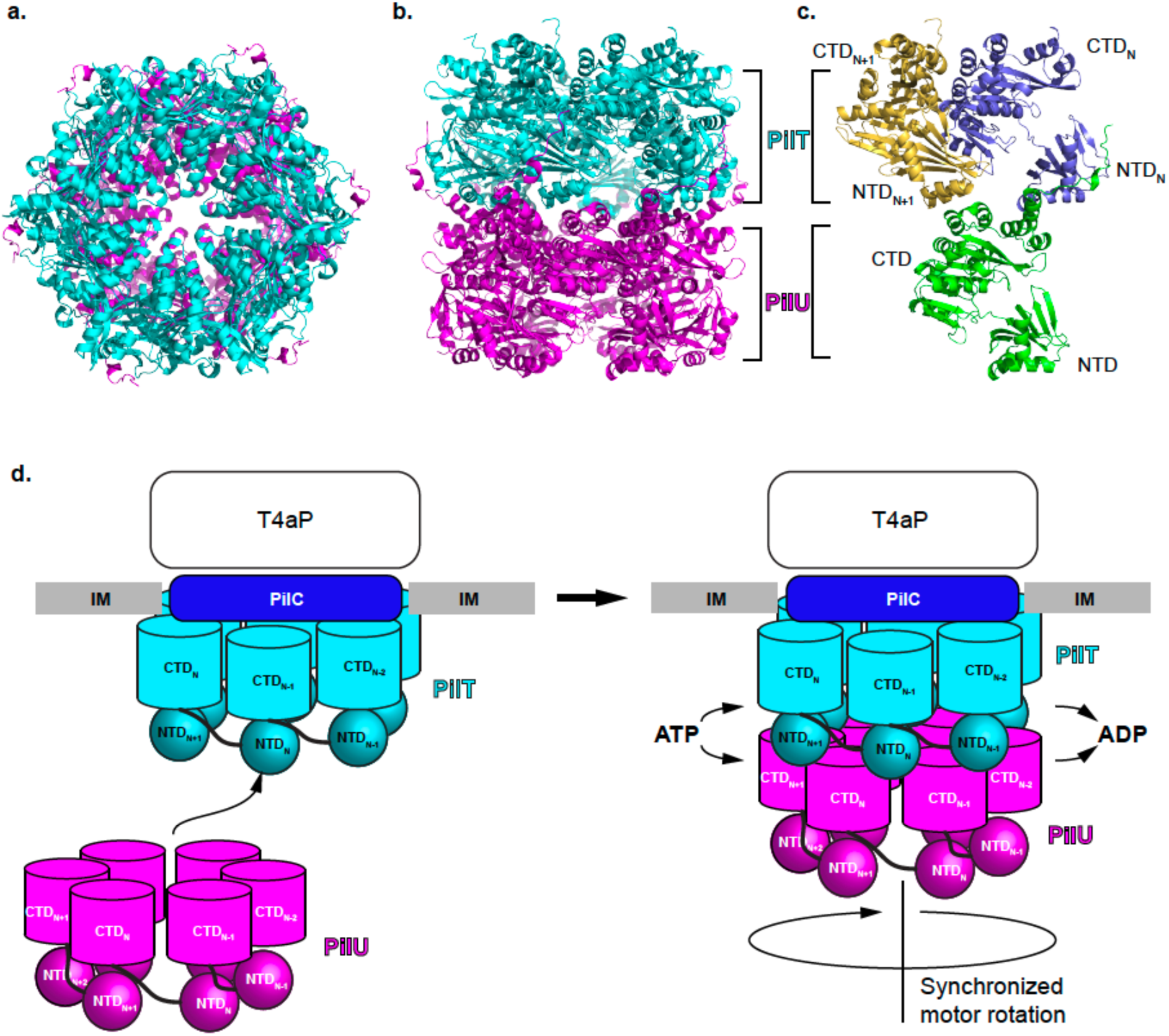
Proposed model of the PilT and PilU functions. a) AlphaFold model of PilT/PilU complex, top view. b) AlphaFold model of PilT/PilU complex, side view. c) AlphaFold model of one PilU subunit interacting with two PilT subunits through the CTD of PilU and NTDs of PilT. d) Proposed mechanism for PilT and PilU recruitment to the T4P machinery.

It is worth noting that direct interaction between PilU and PilC were observed through bacterial adenylate cyclase two-hybrid (BACTH) assay. However, the PilU and PilC interaction is likely too weak for the machinery to function without the presence of PilT in DNA uptake, which is expected because PilU and PilT are adjacent in the operon and co-expressed in *V. cholerae* (Chlebek et al., 2019).

Future studies could be designed to test how PilT and PilU function together during pilus retraction, guided by the available structures. For example, PilU ATPase activity could be measured in the presence of other components of the retraction machinery, such as PilT and PilC, or in the MSHA system with PilT and MshG. In addition, PilU mutations that stabilize a more tightly closed CTD interface could be assessed biochemically to determine their effects on ATP binding and hydrolysis, and structurally to evaluate whether they shift or lock PilU in specific conformational states. Finally, systematic mutagenesis across the PilT and PilU surfaces assisted by AlphaFold models or evolutionary coupling analysis could be performed *in vitro* and *in vivo* to map residues involved in potential PilT/PilU interactions and to validate the functional importance of these interactions during pilus retraction. Teipen *et al* has already performed mutagenesis *in vivo* using predictions from AlphaFold3 and MD simulation and identified a specific salt bridge between K36 in PilT and D366 in PilU that is critical to stabilize the PilT/PilU interaction in *V. cholerae* (Teipen et al., 2025).

## Conclusion

In this study, we determined four cryo-EM and two crystallographic structures of the PilU homohexamer, revealing C1, C2, and C3 symmetries with multiple conformations. No nucleotide was detected at the active site, even with ATP or ADP present in the sample. PilU adopts closed, intermediate, and open conformations with arrangements such as OCOCOC, XOCXOC, COCCOC, COXCOX, XXXXXX and XOCCOC. Transition from closed to open involves loss of CTD_N_-CTD_N+1_ contacts while CTD_N_-NTD_N+1_ interactions persist, stabilizing the hexamer.

PilU resembles its homolog PilT but exhibits less extreme conformational changes. We propose that PilU’s loosely packed closed interfaces underlie its weak ATPase activity, explaining the absence of nucleotide-bound states. PilT may recruit PilU to the pilus retraction machinery, stabilizing a compact interface and enhancing PilU’s ATPase activity.

This system is substantially more dynamic than indicated by structural modeling both by us and by others (Teipen et al., 2025). While modern prediction tools such as AlphaFold are rapidly advancing our understanding of protein structures and interactions, they typically capture a single conformational state. For highly dynamic systems such as PilU, experimental structures remain essential for defining the range of accessible conformations and for elucidating its mechanism.

Together, this report of PilU structures provides six conformational states that will guide mutagenesis, aid in identifying stabilized or nucleotide-bound forms, and clarify PilU’s regulatory role in T4P retraction.

## Acknowledgement

We thank Ankur Dalia for sharing the PilU expression vector and for providing helpful insight at the outset of this project.

This work was supported by HHS/NIH/NIAID contracts HHSN272201700060C (to K.S.), 75N93022C00035 (to K.S. and D.B), R35 GM118108 (to A.M.) and R35 GM145365 (to Z.O.). This research also used resources of the Advanced Photon Source, a U.S. Department of Energy (DOE) Office of Science User Facility operated for the DOE Office of Science by Argonne National Laboratory under Contract No. DE-AC02-06CH11357. Use of the LS-CAT Sector 21 was supported by the Michigan Economic Development Corporation and the Michigan Technology Tri-Corridor (Grant 085P1000817). Access to LS-CAT and use of the Northwestern University Cryo-EM and computational facilities is supported by the Northwestern Structural Biology Facility, which is funded in part by the Robert H. Lurie Comprehensive Cancer Center grant NCI P30CA060553. We thank the Purdue Cryo-EM Facility for the use of the Titan Krios microscope.

The content is solely the responsibility of the authors and does not necessarily represent the official views of the National Institutes of Health. This manuscript is the result of funding in whole or in part by the National Institutes of Health (NIH). It is subject to the NIH Public Access Policy. Through acceptance of this federal funding, NIH has been given a right to make this manuscript publicly available in PubMed Central upon the Official Date of Publication, as defined by NIH.

## Author contributions

Conceptualization: Y.G., S.S., D.B., and K.S.; Methodology: Y.G., S.S., G.M., D.B., and K.S.; Investigation: S.S. performed protein expression, purification, protein crystallization and grid preparation; S.S. and T.K. collected cryo-EM data; Y.G., S.S., T.K., V.T. and A.M. performed single-particle reconstruction; Y.G. validated models and maps; S.S. and G.M. performed X-ray data collection and data analysis; Y.G. performed structure-guided analysis and AlphaFold modeling; Y.G. and Z.O. worked on mechanistic models. Supervision: D.B. and K.S.; Writing - original draft: Y.G.; Writing - review and editing: Y.G., S.S., G.M., N.I., V.T., A.M., D.B., and K.S.; Financial support: A.M., ZO., D.B. and K.S.

---

**Figure S1.**
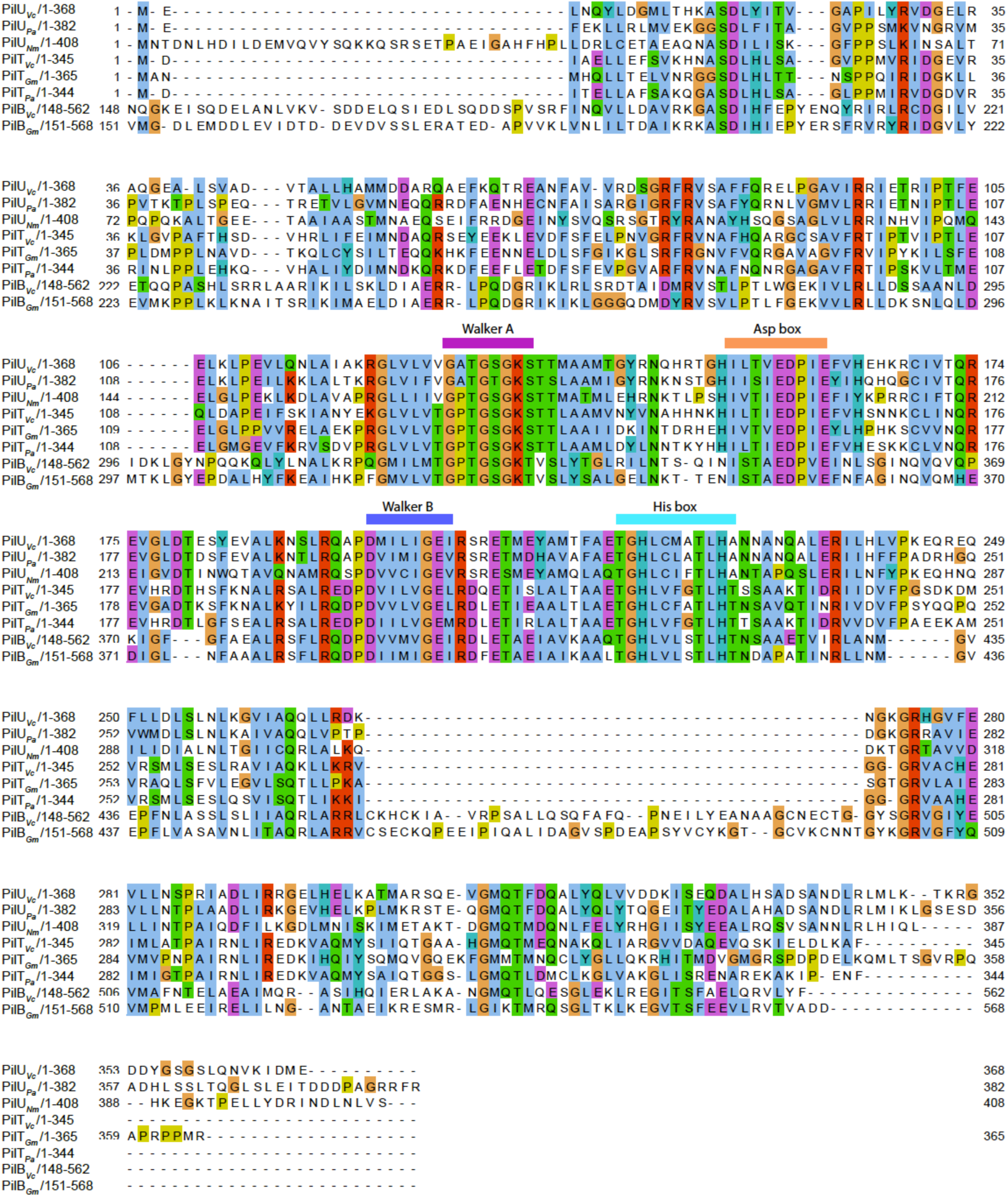
Sequence alignment of PilU, PilT and PilB.

**Figure S2.**
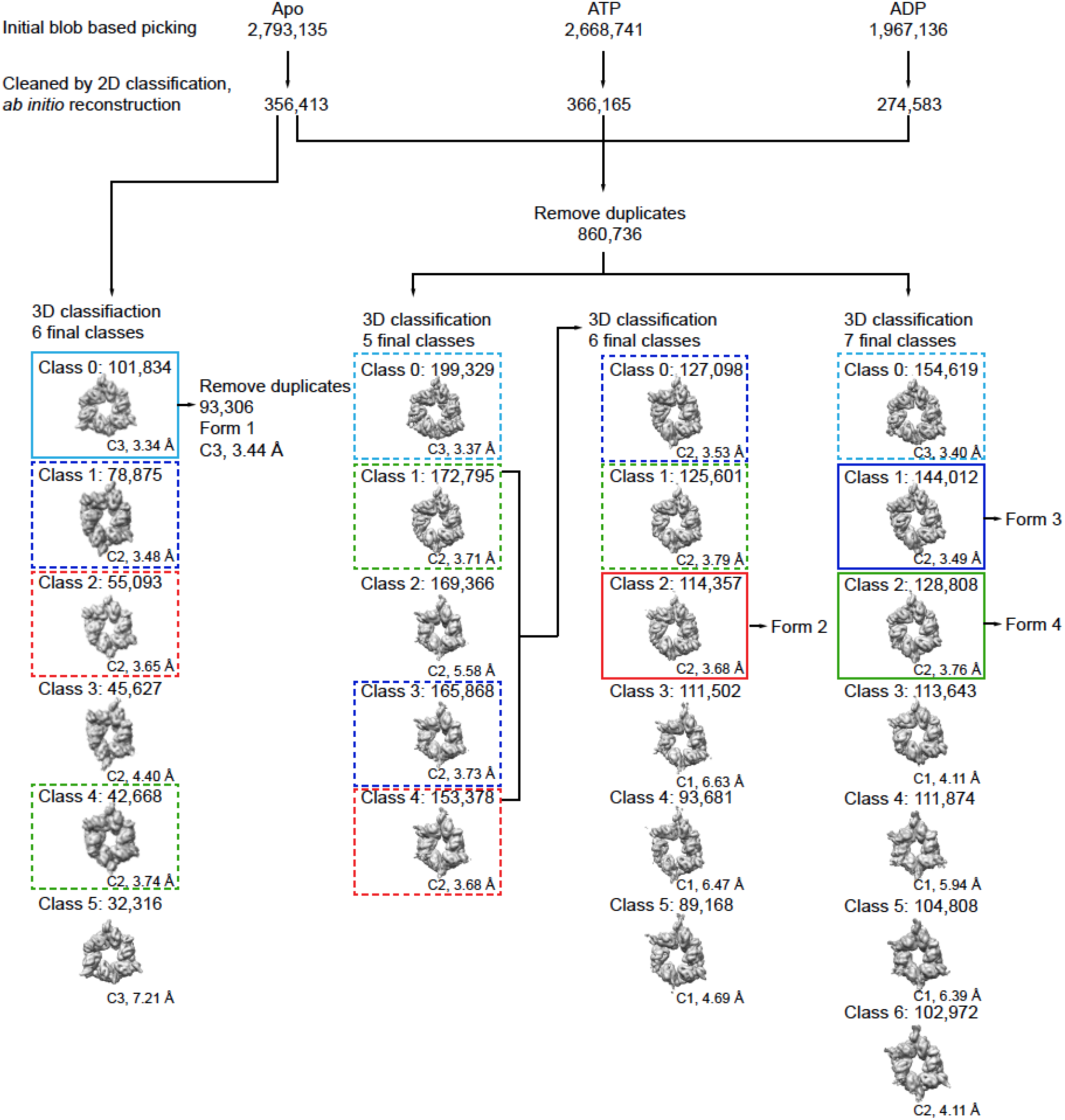
Cryo-EM particle flow. Three datasets (dataset 1: apo, 3,559 movies; dataset 2: in the presence of ATP, 3,788 movies; dataset 3: in the presence of ADP, 3,836 movies) were collected and processed independently to select clean particles. From dataset 1, 3D classification followed by homogeneous refinement yielded PilU form 1 (C3 symmetry, 3.44 Å, cyan box). Qualitative inspection of 2D class averages showed no obvious nucleotide-dependent differences in overall particle shape, apparent symmetry, or distribution of major views; therefore, particles from all three datasets were pooled for further analysis. Multiple rounds of 3D classification and homogeneous refinement of the combined particle set resolved PilU form 2 (C2 symmetry, 3.68 Å, red box), form 3 (C2 symmetry, 3.49 Å, blue box), and form 4 (C2 symmetry, 3.76 Å, green box). Similar conformations corresponding to PilU forms 1-4 were observed across multiple datasets and 3D classification jobs (cyan dashed box: classes similar to form 1; red dashed box: classes similar to form 2; blue dashed box: classes similar to form 3; green dashed box: classes similar to form 4), further indicate that the addition of ATP and ADP did not substantially change the overall PilU conformations, even if the relative population of particles in each class may differ. For clarity, only the highest-quality reconstruction of each form is used for atomic model building and subsequent analysis. 3D reconstructions with nominal resolutions worse than ∼4 Å, which may represent minor conformations or other poorly resolved classes, lacked interpretable structural features and were therefore not assigned as discrete PilU forms and were not further analyzed. Differences in particle numbers across similar reconstructions from individual classification jobs likely reflect the stochastic nature of 3D classification in the presence of continuous heterogeneity and its sensitivity to parameter choices, rather than significant shifts in conformational populations.

**Figure S3.**
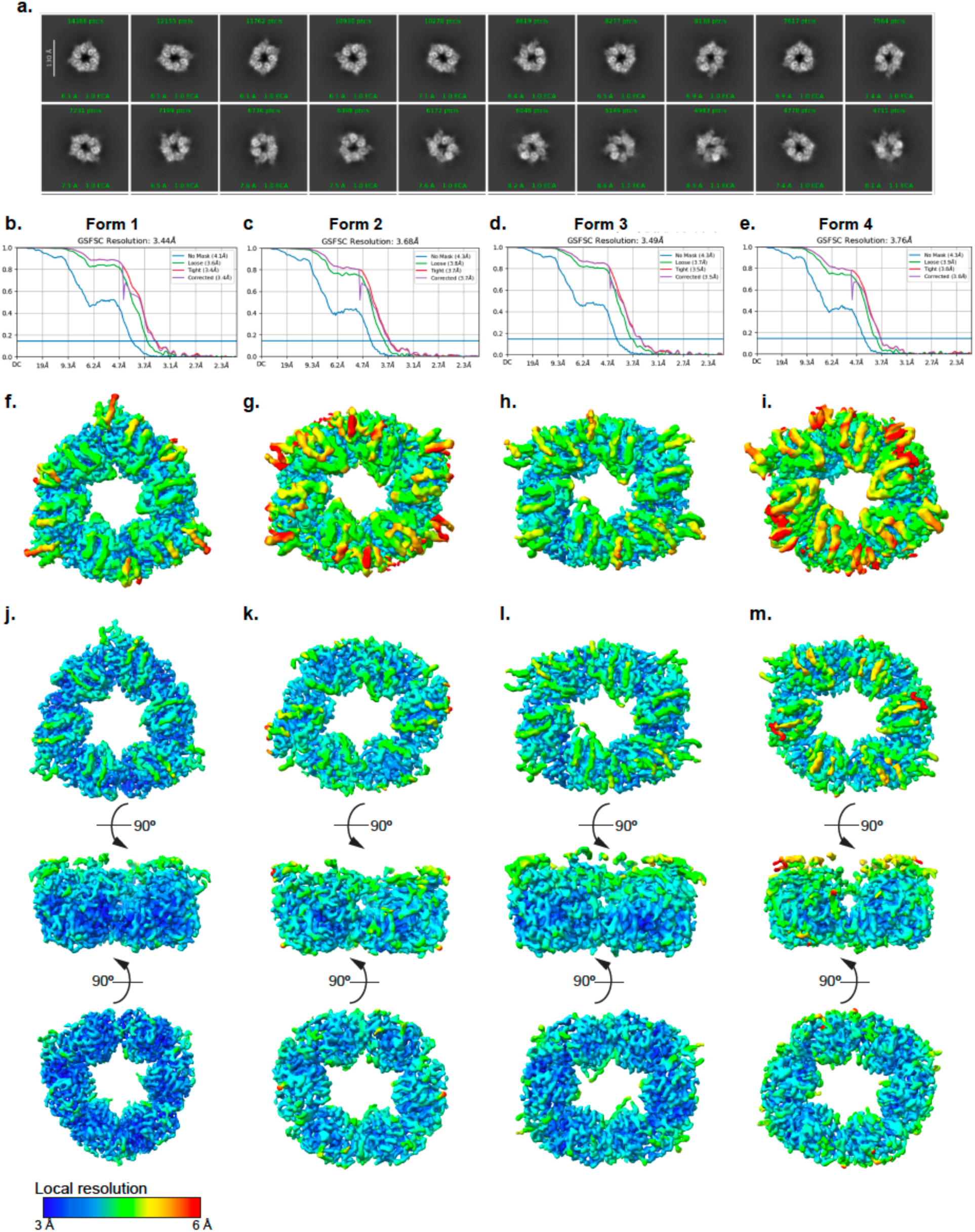
2D classes and electron density maps. a) 2D classes of PilU show different conformations of hexameric rings. b-e) Global FSC of forms 1-4. f-i) Low contour local resolution maps of forms 1-4. j-m) High contour local resolution maps of forms 1-4.

**Figure S4.**
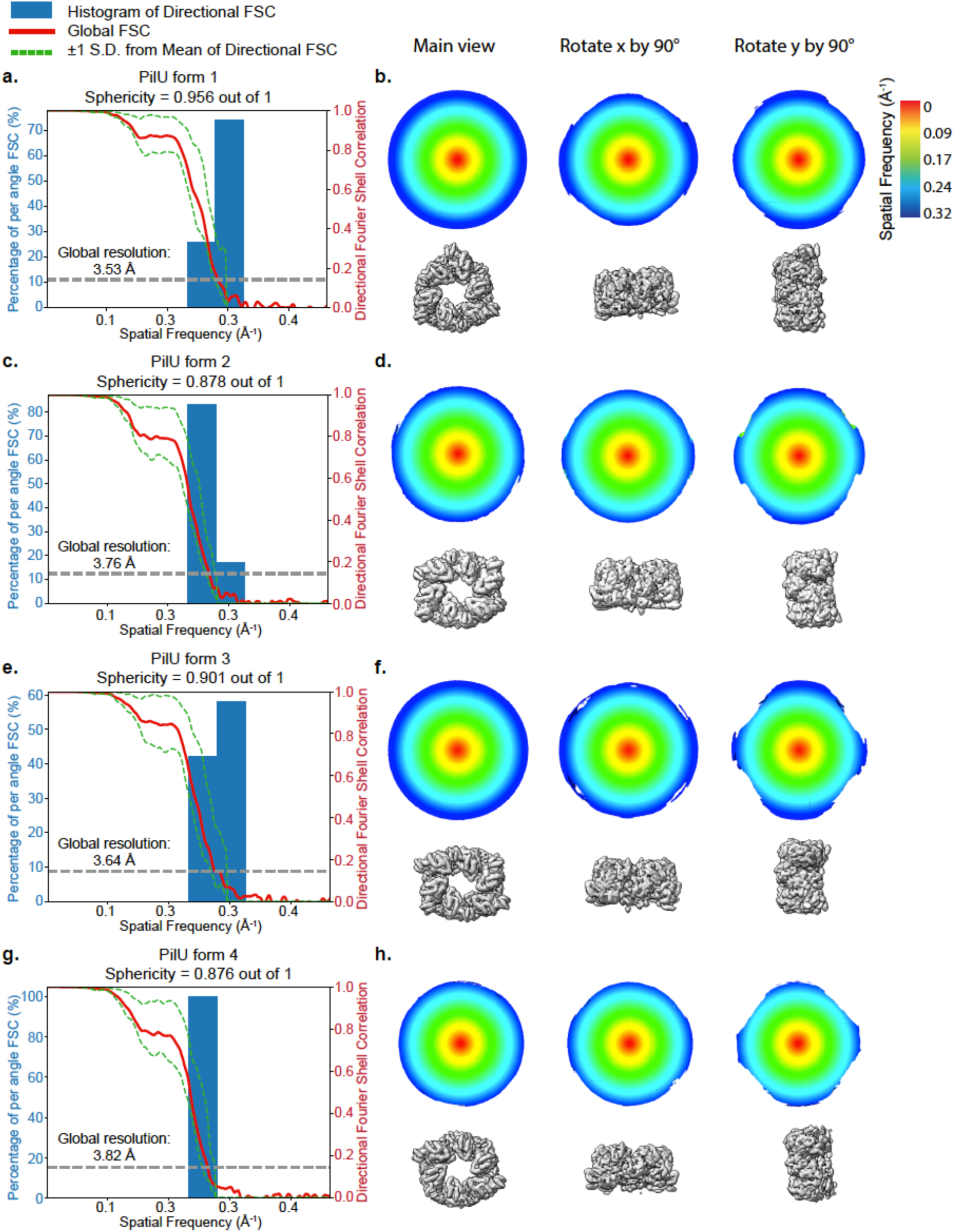
Resolution assessment of PilU forms 1-4 by directional FSC. Global and directional Fourier shell correlation curves calculated between half maps with masks used in the final 3D refinement are shown for PilU forms 1-4 (global FSC: solid red line; ±1 standard deviation from mean of directional FSC: dashed green line). Global resolutions estimated by 3DFSC are slightly lower (0.06-0.15 Å) than those estimated by CryoSPARC. Directional resolution anisotropy was assessed by 3D FSC analysis, visualized as FSC volumes rotated by 90° about the x and y axes. The narrow distribution of the per angle FSC histogram and the near-spherical 3D FSC distributions indicate minimal preferred particle orientation and limited resolution anisotropy for all four reconstructions.

**Figure S5.**
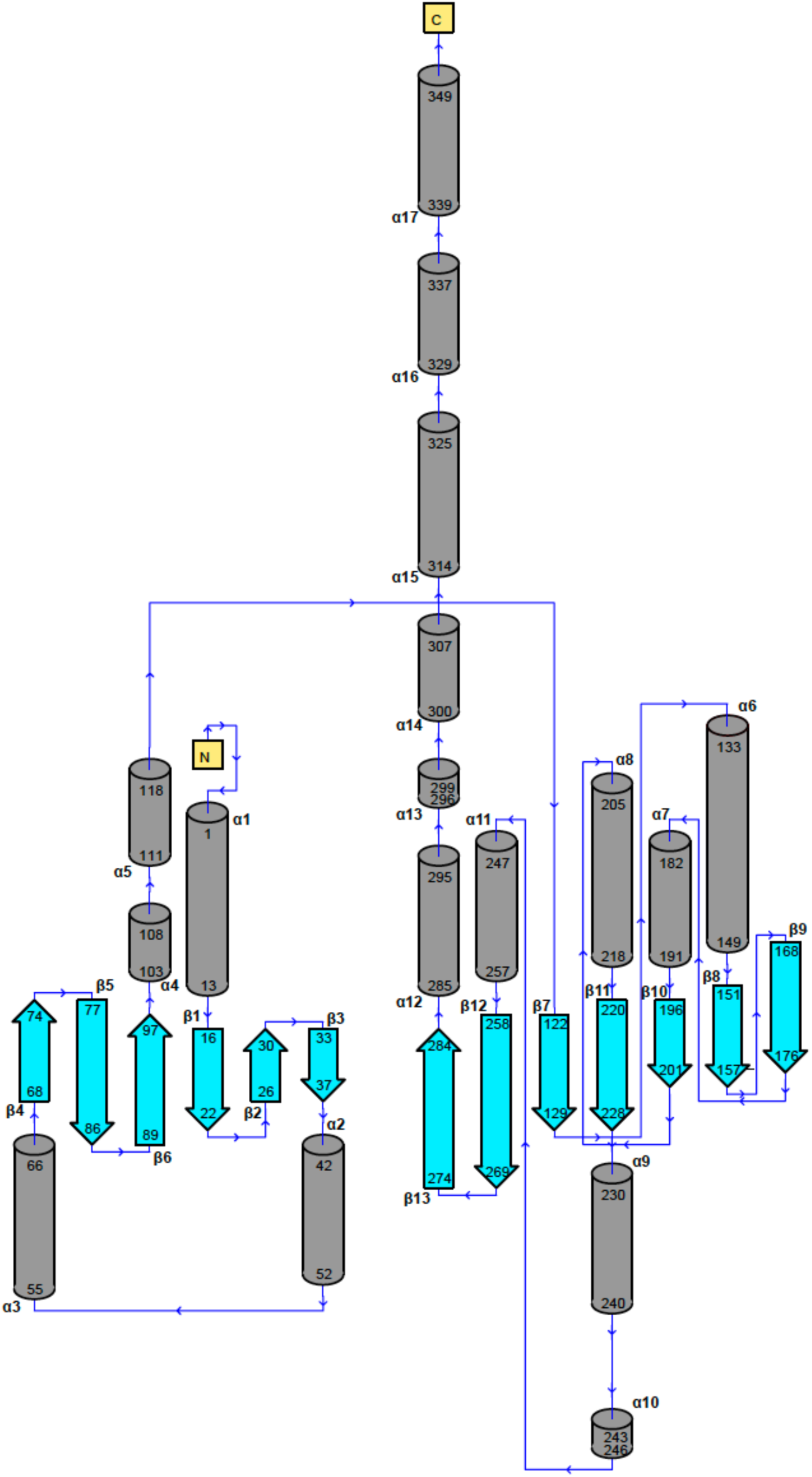
Secondary structure of PilU subunit.

**Figure S6.**
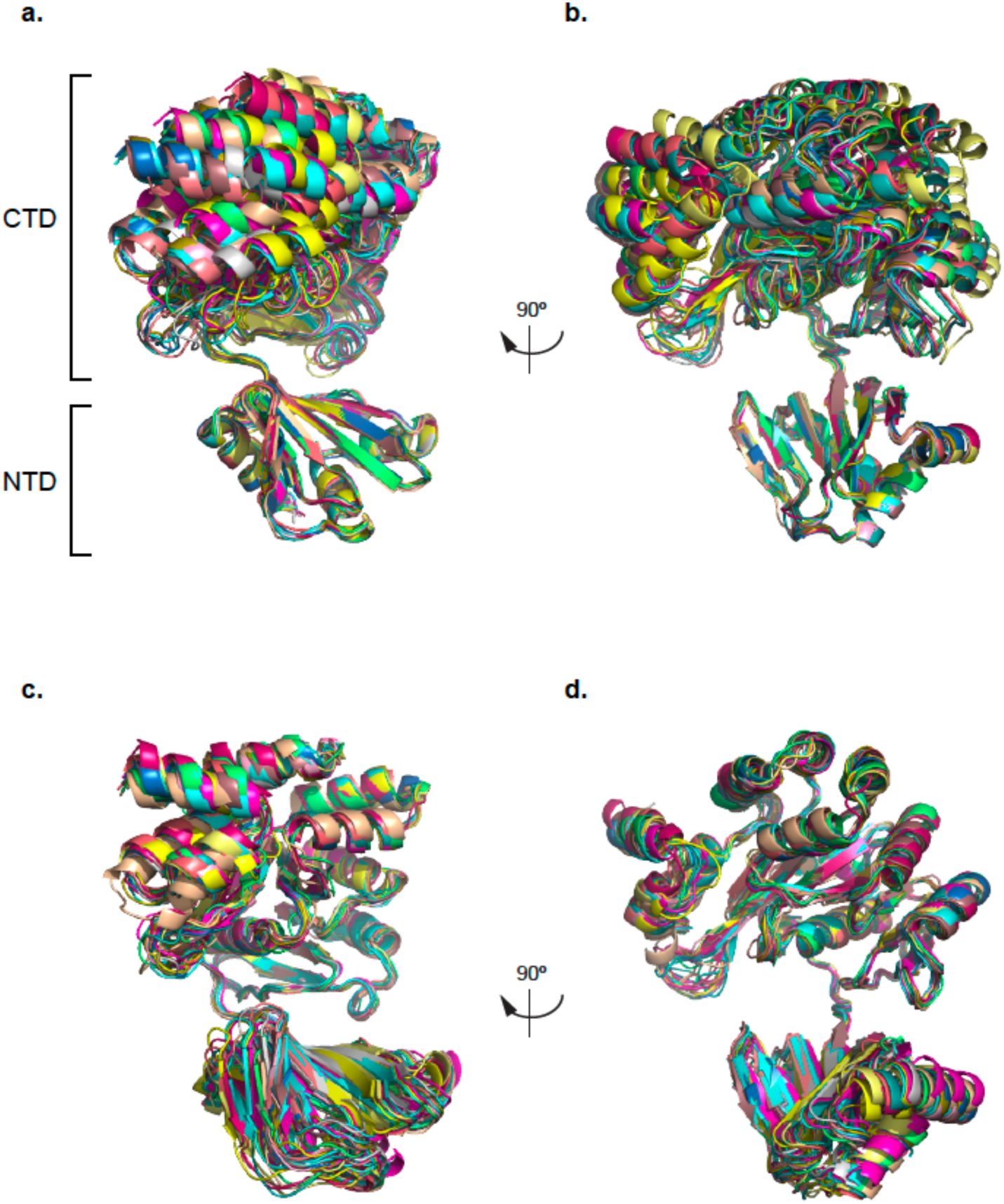
Superimposed PilU subunits. a, b) PilU subunits superimposed on NTD. c, d) PilU subunits superimposed on CTD.

**Figure S7.**
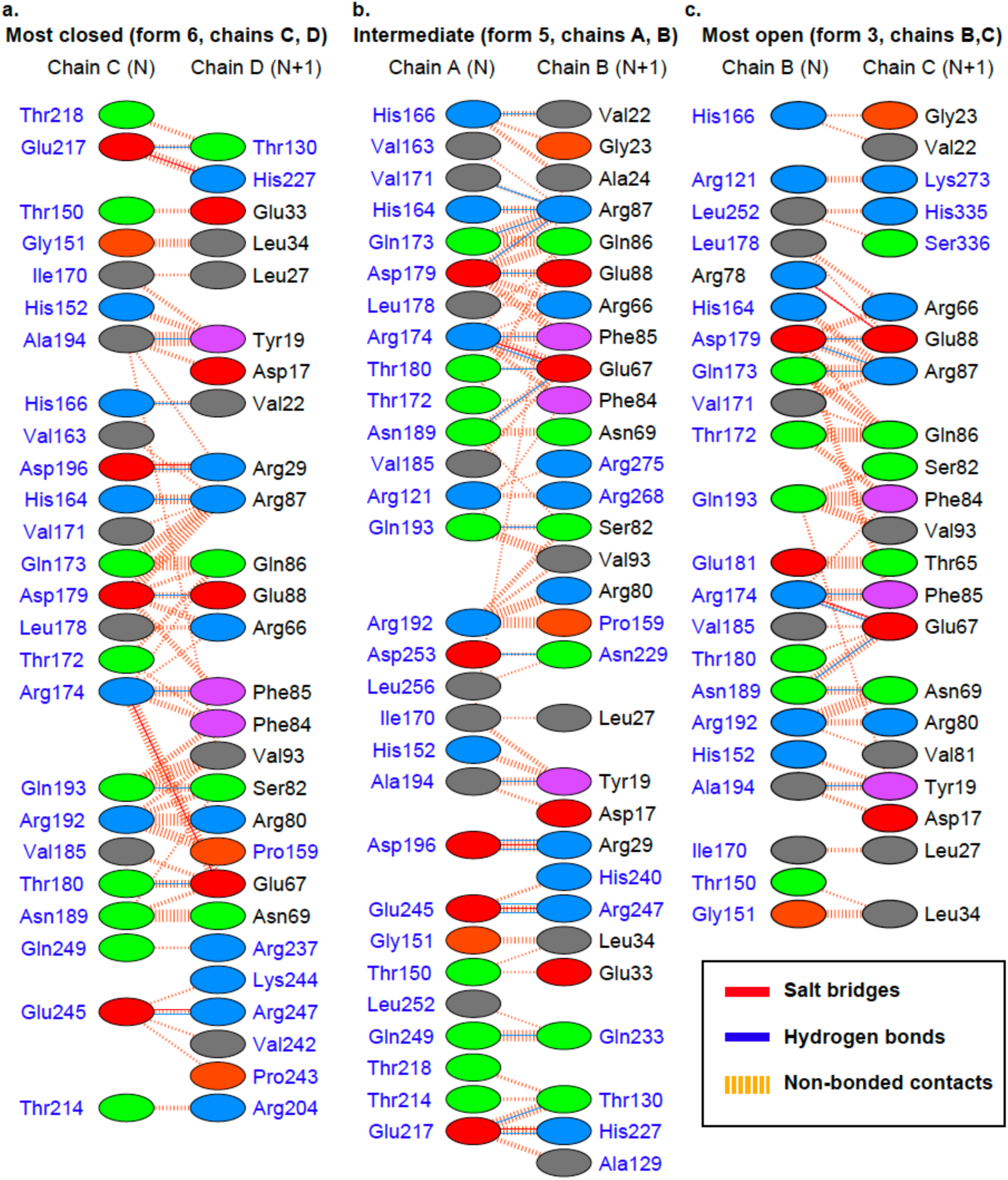
Contacts between PilU subunit interface. Residues in the NTD are shown in black text, and residues in the CTD are shown in blue text. From the closed to the open state, contacts between CTD_N_ and CTD_N+1_ are significantly reduced.

**Figure S8.**
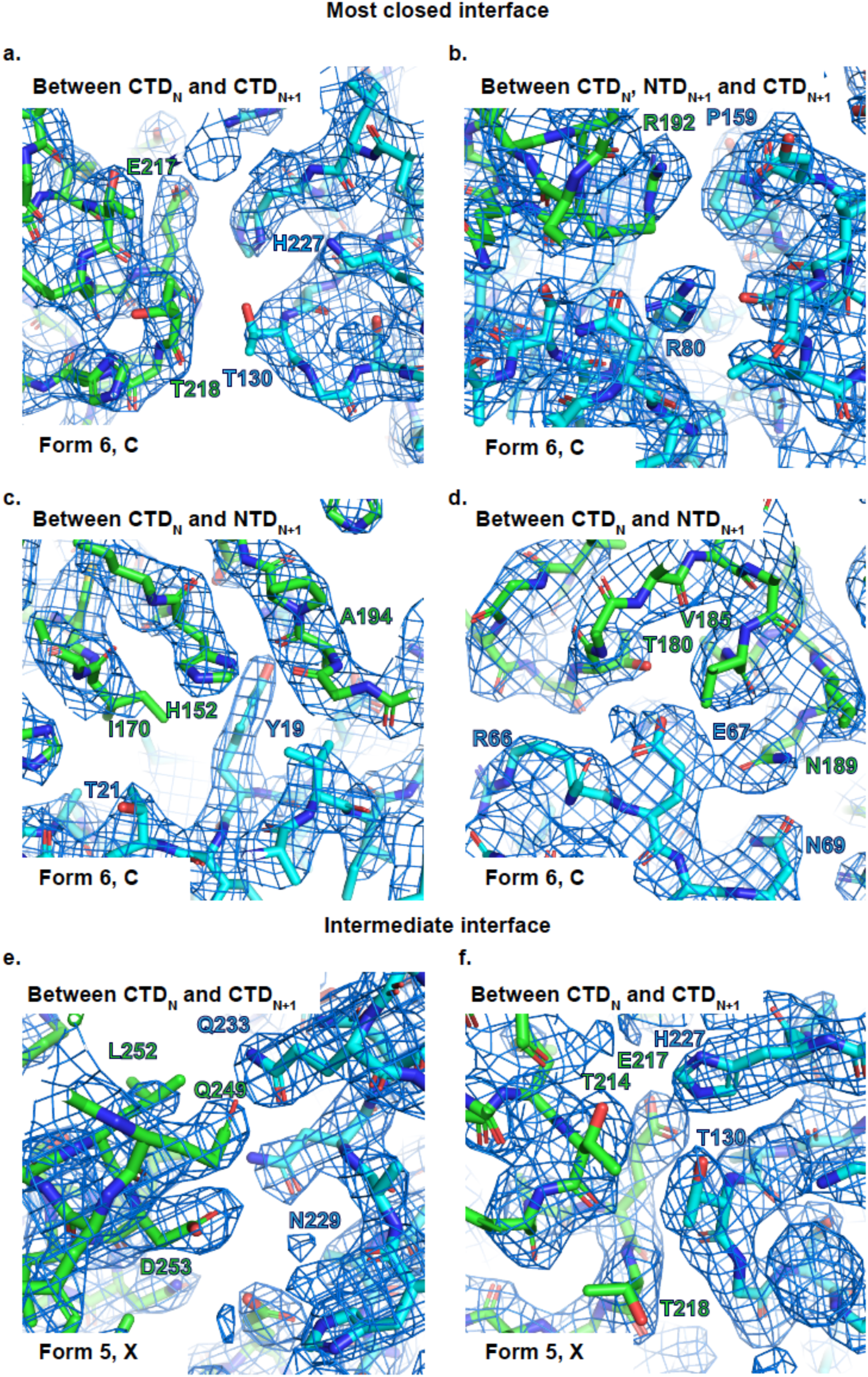
Density at PilU subunit interface. Chain_N_ is shown in green, and chain_N+1_ is shown in cyan.

